# Next generation infection prevention clothing: Non-woven Fabrics Coated with Cranberry Extracts Capable of Inactivating Enveloped Viruses such as SARS-CoV-2 and Multidrug-resistant Bacteria

**DOI:** 10.1101/2021.08.14.456330

**Authors:** Alberto Tuñón-Molina, Alba Cano-Vicent, Miguel Martí, Yukiko Muramoto, Takeshi Noda, Kazuo Takayama, Ángel Serrano-Aroca

**Affiliations:** Biomaterials and Bioengineering Lab, Centro de Investigación Traslacional San Alberto Magno, Universidad Católica de Valencia San Vicente Mártir, c/Guillem de Castro 94, Valencia 46001, Spain; Laboratory of Ultrastructural Virology, Institute for Frontier Life and Medical Sciences, Kyoto University, Kyoto 606-8507, Japan; Center for iPS Cell Research and Application, Kyoto University, Kyoto 606-8397, Japan

**Keywords:** SARS-CoV-2, non-woven fabric, cranberry extract, antimicrobial activity, infection prevention clothing

## Abstract

The Coronavirus Disease (COVID-19) pandemic is demanding rapid action of the authorities and scientific community in order to find new antimicrobial solutions that could inactivate the pathogen SARS-CoV-2 that causes this disease. Gram-positive bacteria contribute to severe pneumonia associated with COVID-19, and their resistance to antibiotics is increasing at an alarming rate. In this regard, non-woven fabrics are currently used for the fabrication of infection prevention clothing such as face masks, caps, scrubs, shirts, trousers, disposable gowns, overalls, hoods, aprons and shoe covers as protective tools against viral and bacterial infections. However, these non-woven fabrics are made of materials that do not possess antimicrobial activity. Thus, we have developed here non-woven fabrics with antimicrobial coatings of cranberry extracts capable of inactivating enveloped viruses such as SARS-CoV-2 and the phage phi 6, and two multidrug-resistant bacteria: the methicillin-resistant *Staphylococcus aureus* and *Staphylococcus epidermidis*. The non-toxicity of these advanced technology was ensured using a *Caenorhabditis elegans in vivo* model. These results open up a new prevention path using natural and biodegradable compounds for the fabrication of infection prevention clothing in the current COVID-19 and future pandemics.

## 1. Introduction

Severe Acute Respiratory Syndrome Coronavirus 2 (SARS-CoV-2) is the third human coronavirus ^1–4^ that is much more contagious than SARS-CoV and MERS-CoV ^5–12^. This rapid transmission rate has provoked the current Coronavirus Disease (COVID-19) pandemic. SARS-CoV-2 is a highly pathogenic enveloped positive-sense single-stranded RNA virus ^13–15^ that belongs to the Baltimore group IV ^16^. This global life-threatening situation needs the development of new antimicrobial approaches that could treat or prevent COVID-19 infections^17–22^. In this regard, non-woven fabrics are currently used for the fabrication of infection prevention clothing such as face masks, caps, scrubs, shirts, trousers, disposable gowns, overalls, hoods, aprons, and shoe covers. These infection prevention tools are needed specially in hospitals during surgical operations, in microbiological and biomedical biosafety laboratories and also, in the case of face masks, by most citizens as a demonstrated prevention tool in the current COVID-19 pandemic. Nevertheless, these infection prevention clothing are produced with materials that do not possess antimicrobial properties. Many disinfectants such as household bleach, hand soap solution, ethanol, povidone-iodine, chloroxylenol, chlorhexidine and benzalkonium chloride have shown potent antiviral activity against SARS-CoV-2 so far ^23^. Thus, we have treated non-woven fabrics with benzalkonium chloride to obtained composite fabrics capable of inactivating SARS-CoV-2 and multidrug-resistant bacteria ^24^. However, other natural and biodegradable compounds such as cranberry extracts have not been tested yet. Cranberry extracts have shown antiviral activity against other enveloped viruses such as the herpes simplex virus type 1 (HSV-1) and type 2 (HSV-2) due to the presence of antimicrobial A-type proanthocyanidins (PACs) that provoke alterations of their envelope glycoproteins. However, although HSV-1 and HSV-2 belong to a different Baltimore group I ^16^ than SARS-CoV-2 because they are double-stranded DNA viruses, they are also enveloped viruses like SARS-CoV-2. A cranberry extract has also exhibit antiviral activity against influenza virus (IFV) ^25^. IFV is a negative-sense single-stranded RNA virus that belongs to the Baltimore group V ^16^. However, it is also enveloped like HSV-1, HSV-2. Therefore, since it seems that the PACs present in cranberry extracts effectively interact with the envelope glycoproteins achieving viral inhibition, we hypothesize here that a commercial non-woven fabric treated with two different commercial extracts produced with different cranberries will show antiviral activity against the enveloped SARS-CoV-2 and phi 6 viruses. The phage phi 6 is also an enveloped double-stranded RNA virus (group III of the Baltimore classification ^16^) that can be used as biosafe viral model of SARS-CoV-2 and other enveloped viruses such as influenza due to safety reasons ^24^. Furthermore, atypical viral pneumonia is associated with SARS-CoV-2 infection ^2,26^ that can increase its risk by co-infection with Gram-positive bacteria ^27–30^, including clinically relevant antibiotic-resistant strains. Additionally, bacterial resistance to pneumonia treatments is increasing at an alarming rate ^31,32^. Since the PACs present in cranberry extracts are well-known for their antibacterial properties against Gram-negative *Escherichia coli* ^33^ and antifungal activity against *Candida albicans* ^34^, we hypothesize here also that the two non-woven fabrics dip-coated with cranberry extracts will show also antibacterial activity against two Gram-positive multidrug-resistant bacteria, the methichillin *Staphylococcus aureus* (MRSA) and *Staphylococcus epidermidis* (MRSE).

## 2. Materials and Methods

### 2.1. Fabric preparation

Cranberry extracts with a concentration of 10% *w/v* were prepared using ethanol as extracting solvent and two different types of commercial cranberry powders: VITAFAIR (*Vaccinium macrocarpon*, Germany) and NUTRIBIOLITE (Uritractin, *Vaccinium macrocarpon*, Spain). According to the VITAFAIR and NUTRIBIOLITE manufacturers, the cranberry powders have a proanthocyanidins content of 25 and 30% *w/w*, respectively. Thus, 10 grams of cranberry powder was mixed with 100 mL of absolute ethanol (≥99.8%, VWR chemicals, AnalaR Normapur) under magnetic stirring for 30 minutes at 24±1°C sealed with parafilm. After that, the extract was left overnight to decant the solid phase. After 12 hours, the supernatant was centrifugated at 10000 r.p.m (Centrifuge Heraeus Megafuge 16R, Thermo Scientific) for 30 minutes to ensure complete phase separation of the liquid cranberry extract from the solid phase. After centrifugation, extract was left overnight to decant possible solid phase remaining. The day after, supernatant was removed with a 25mL serological pipette (LABCLINICS) and filtered with a non-sterile 0.2 μm filter. After filtering, extract was centrifuged again at 10000 rpm for 30 minutes to ensure complete solid phase separation from the liquid phase. Thus, commercial non-woven spunlace fabric from NV EVOLUTIA (Valencia, Spain) was prepared in the form of discs of approximately 10 mm in diameter. After that, they were treated with the two different cranberry extracts by the dip-coating method ^35^. Thus, the non-woven fabrics were immersed in the cranberry extracts (10% *w/v*) solutions for 30 minutes at 24±1ºC sealed with parafilm. After that, the prepared samples were dried at 60ºC for 48 hours to solidify the physically absorbed cranberry extract to form the coating and ensure complete evaporation of the ethanol phase. The non-woven fabrics treated with VITAFAIR and NUTRIBIOLITE cranberry extracts will be named hereafter as E10V and E10N, respectively. The discs were sterilized by ultraviolet radiation for one hour per side. Discs prepared from the non-woven fabrics treated with only the absolute ethanol (≥99.8%, VWR chemicals, AnalaR Normapur) solvent under magnetic stirring for 30 minutes at 24±1°C (Control S) were prepared as reference material. All these experiments were performed in triplicate to ensure reproducible results.

### 2.2. Fabric morphology

The morphology of the non-woven fabrics treated with and without cranberry extracts was observed by optical microscopy (Motic BA410E) and was photographed at 10x and 40x magnifications with the Moticam 580 5.0MP. The images were processed by the Motic Images Plus 3.0 software. Macroscopic photographs of the fabrics were also performed with a 64MP Xiaomi camera with a Sony IMX682 sensor and a f/1.89 opening.

### 2.3. Antiviral test against SARS-CoV-2

The SARS-CoV-2 (SARS-CoV-2/Hu/DP/Kng/19-027) was provided by Dr. Tomohiko Takasaki and Dr. Jun-Ichi Sakuragi (Kanagawa Prefectural Institute of Public Health). A volume of 100 μL of a SARS-CoV-2 suspension in PBS was added to each disc at a titer dose of 5.0×10^6^ median tissue culture infectious dose (TCID50)/disc, and then incubated for 1 minute at room temperature. 900 μl PBS were added to each disc, and then vortexed for 5 minutes. After that, each disc was vortexed for 5 minutes. Briefly, TMPRSS2/Vero cells (JCRB1818, JCRB Cell Bank) were cultured with the Minimum Essential Media (MEM, Sigma-Aldrich) containing 5% fetal bovine serum (FBS), 1% penicillin/streptomycin (P/S) on the 96-well plates (Thermo Fisher Scientific). Samples were serially diluted 10-fold from 10^−1^ to 10^−8^ in the MEM containing 5% FBS and 1% P/S. Dilutions were placed onto the TMPRSS2/Vero cells in triplicate and incubated at 37°C for 96 hours. Cytopathic effects were evaluated under a microscope. TCID50/mL were calculated using the Reed-Muench method. The SARS-CoV-2 infection experiments were conducted at a Biosafety Level 3 laboratory at Kyoto University.

### 2.4. Antiviral test against phi6

*Pseudomonas syringae* (DSM 21482) is the host of the phage phi 6. This Gram-negative bacterium was purchased from the Leibniz Institute DSMZ–German Collection of Microorganisms and Cell cultures GmbH (Braunschweig, Germany). This microorganism was cultured in solid tryptic soy agar (TSA, Liofilchem) and, after that, in liquid tryptic soy broth (TSB, Liofilchem) at a speed of 120 r.p.m. at 25 °C. The Leibniz Institute DSMZ–German Collection of Microorganisms and Cell Cultures GmbH specifications were followed to propagate the phage phi 6 (DSM 21518). The antiviral assay was performed with a dispersion of 50 μL of TSB with phages placed onto each sample at a titer of approximately 1 × 10 ^6^ plaque-forming units per mL (PFU/mL) and incubated for 1 minute. A falcon tube was used to place each disc with 10 mL TSB to be sonicated and vortexed for 5 and 1 minute, respectively at ambient temperature (24 ±1°C). Phage titration by serial dilutions of each falcon sample was performed and 100 μL of each phage dilution was mixed with 100 μL of the bacterial host at OD_600nm_=0.5. The infective capacity of the phage phi 6 was analyzed based on the double-layer method ^36^. Thus, a volume of 4 mL of top agar (TSB + 0.75% bacteriological agar, Scharlau) and 5 mM calcium chloride (Sigma-Aldrich) were added to the phage dispersion mixed with the bacteria, and then poured on TSA plates for incubation for 18–24 hours in a refrigerated oven at 25 °C. Phage titers in PFU/mL of each sample were compared with control (CONTROL), which consisted of 50 μL of phage added directly to the bacterial culture without being in contact with any type of disc and without the sonication/vortexing treatment. The antiviral activity of the discs coated with cranberry extract or not was determined at 1 minute of contact with the phage phi 6 in log reductions of titers. It was made sure that the sonication/vortexing treatment did not affect the infectious activity of the phage phi 6 and that the residual disinfectants of the titrated samples did not interfere with the titration process. These antiviral tests were performed in triplicate during two different days (*n* = 6) to achieve reproducibility.

### 2.5. Antibacterial Tests

Lawns of MRSA, COL ^37^, and MRSE, RP62A ^38^, were used to perform the antibacterial assays by the agar disc diffusion tests ^39,40^ at a concentration of approximately 1.5 × 10^8^ CFU/mL in tryptic soy broth, and then cultivated on trypticase soy agar plates. Incubation was performed aerobically at 37 °C for 24 h with the sterilized samples treated with (E10V and E10N) and without cranberry extract (Control S) placed upon them. The inhibition zone (or halo) was normalized according to Equation (1) ^39^.

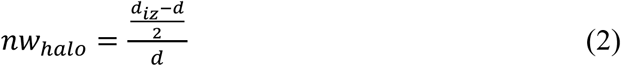

In this equation, *nw*_*halo*_ expresses the normalized width value of the antibacterial inhibition zone, *d*_*iz*_ indicates the diameter of the inhibition zone and the term *d* represents the diameter of the disc. The diameter of the disc was measured by image software analysis (Image J, Wayne Rasband (NIH), USA). These antibacterial assays were performed in triplicate during two different days (*n* = 6) to provide reproducible results.

### 2.6. *In vivo* toxicity tests

*In vivo* toxicity was studied in the *Caenorhabditis elegans* model. Extractions from the non-woven fabrics with (Control S) and without (E10V and E10N) cranberry extract treatment were accomplished following the ISO-10993 standard recommendations. Thus, spunlace fabrics treated with and without cranberry extracts were subjected to sterilization under ultraviolet light (1 hour per side). A 6-well plate was used to place every piece of fabric into a well with 2 mL of potassium medium (2.36 g potassium chloride, 3 g sodium chloride in 1 L distilled water, autoclaved). A volume ratio of 0.1g/mL was selected according to the ISO-10993 that recommends this rate for irregular porous materials of low density such as textiles. After incubating for 24 hours at 25ºC, extracts were collected in 1.5 mL eppendorf tubes. The worms were maintained and propagated on OP50 *E. coli* seeded nematode growth medium (NGM) prepared according to Stiernagle, T. ^41^ at 25ºC. N2 strain was used in these experiments provided by the *Caenorhabditis* Genetics Center (CGC, Minneapolis, USA). The worms and eggs were washed off NGM plates using 5 mL of distilled water and collected in 15 mL falcon tubes to prepare an L1 stage-synchronized *C. elegans* population. Tubes were centrifuged at 1300 rpm for 3 minutes and supernatant was moved away. *C. elegans*’ pellet was resuspended in 100 μL of dH_2_O and transferred to eppendorf tubes adding 700 μL of a 5% bleaching solution. This mixture was incubated for 15 minutes while vortexing every 2 minutes. After last vortexing procedure, eppendorf tubes were centrifuged at 700 g for 3 minutes. Supernatant was moved away, and pellet was washed in 800 μL of dH_2_O. This step was carried out two more times. After last washing step, pellet was resuspended in 100 μL of dH_2_O and transferred to NGM plates seeded with 100 μL of an OP50 *E. coli* culture. Eggs were incubated for 72 hours at 25ºC. Centrifugation at 1300 rpm for 3 minutes was performed to pellet the L1 staged populations and subsequent resuspension in 3 mL of potassium medium was performed. A 48-well plate was used to prepared wells with 62.5 μL of a 1:250 suspension of cholesterol (5 mg/mL in ethyl alcohol) in sterile potassium medium, 62.5 μL of a 50X concentrated *E. coli* OP50 culture with an OD of 0.9, pelleted by centrifugation (4000 rpm for 10 minutes) and resuspended in potassium medium, 115 μL of potassium medium and 250 μL of the pertinent cranberries extract. A volume of K medium containing 50-100 worms was then added. 48-well plates were sealed with parafilm and place in an orbital shaker at 25ºC and 120 rpm for 24 hours. In order to determine the survival rate of *C. elegans*, the volume of each well of the 48-well plates was divided in 10 drops of 50 μL and placed under the microscope (Motic BA410E including Moticam 580 5.0MP) in order to count the number of living *C. elegans* and deceased ones. The growth in the medium and in bleach was used a positive and negative control, respectively. To analyze reproduction, three worms were placed into a new OP50 seeded NGM plate and allowed to lay eggs for 48 hours. Then, eggs were counted by using the microscope. Growth was assessed in heat-killed samples by measurement the body length in a picture taken under the microscope with Motic Images Plus 3.0 software. Five replicates were conducted for this assay.

### 2.7. Statistical Analysis

The ANOVA statistical analysis followed by Tukey’s post hoc test (* *p* > 0.05, *** *p* > 0.001) was performed using the GraphPad Prism 6 software (GraphPad Software Inc., USA).

## 3. Results and discussion

### 3.1. Fabric morphology

The optical microscopy images at two magnifications and the macroscopic photographs of the non-woven fabrics treated with and without cranberry extracts are shown in Figure 1.

**Figure 1.**
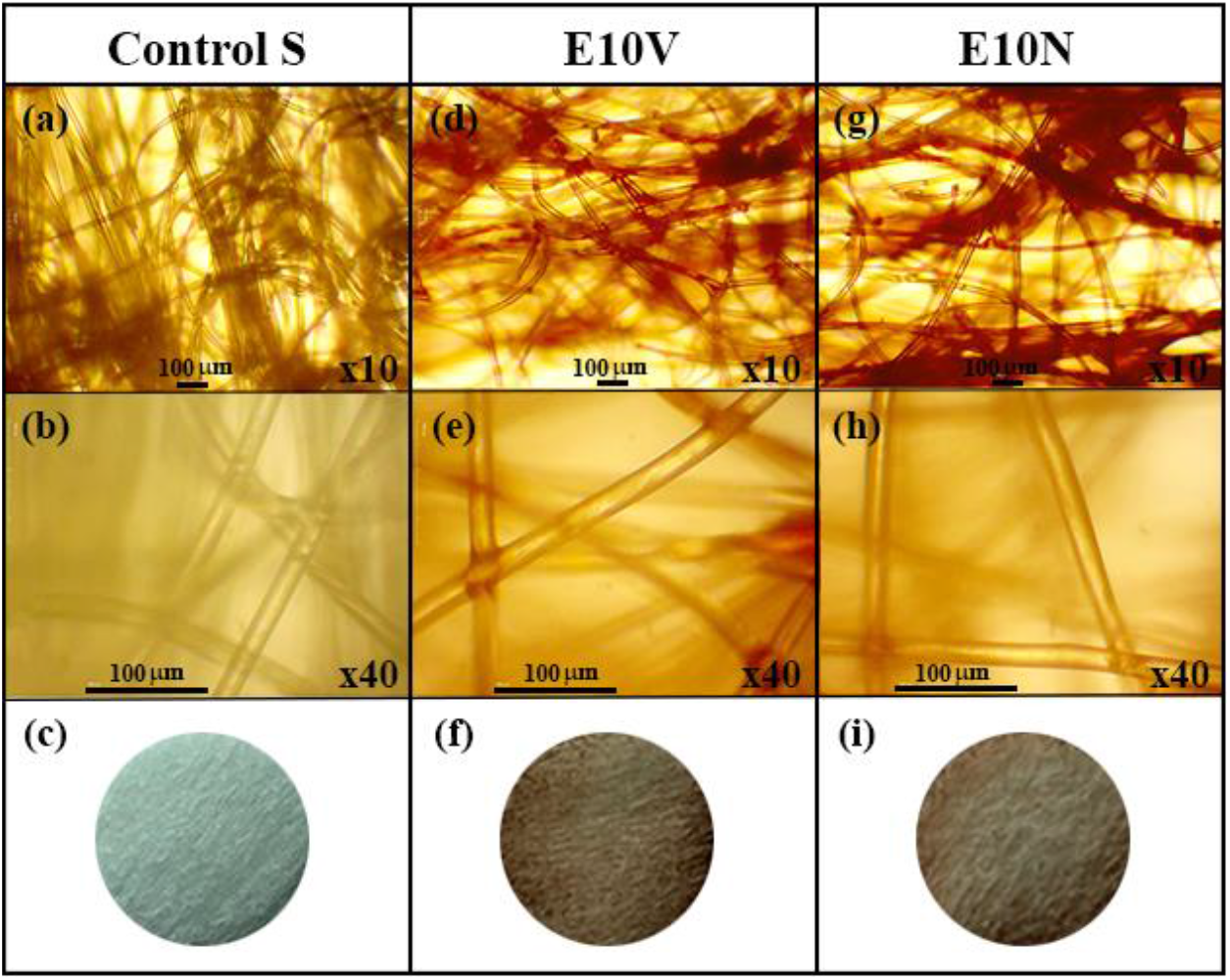
Optical microscopy images at two magnifications (x10 and x40) and the macroscopic photographs of the non-woven fabrics before (a,b,c) and after the treatment with VITAFAIR (E10V) (d,e,f) or NUTRIBIOLITE (E10N) (g,h,i).

Both optical microscopy images and macroscopic photographs show that only a very thin reddish coating of cranberry extract is form onto the fibers of the non-woven fabrics. Thus, no change of porosity or fiber arrangement is observed, which suggests no change of breathability or bacterial filtration efficiency required for certain infection prevention clothing applications.

### 3.2. Antiviral results

The results achieved with SARS-CoV-2 after 1 minute of contact with the non-woven fabric treated with cranberry extracts and the controls are shown in Figure 2.

**Figure 2.**
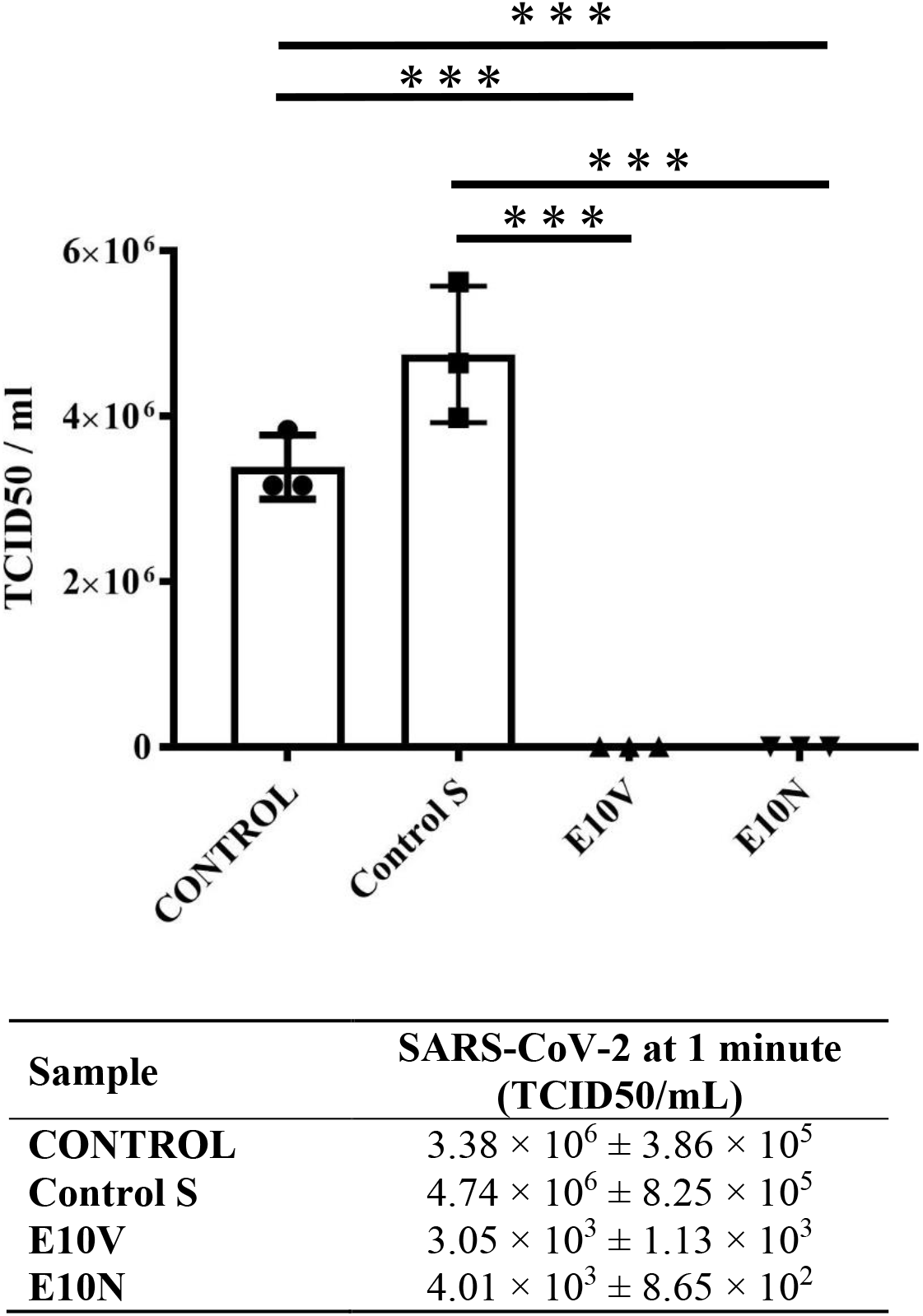
Reduction of infectious titers of SARS-CoV-2 after 1 minute of contact was evaluated by the TCID50/mL method. Control, untreated non-woven fabric (Control S), and non-woven fabric with biofunctional coating of cranberry extracts of VITAFAIR (E10V) or NUTRIBIOLITE (E10N) at 1 minute of viral contact. A dot, square and triangle plot is a data set based on the value of each point. ^***^ *p* > 0.001; ns, not significant. Data are represented as means ± SD.

Therefore, the non-woven fabric E10V and E10N showed strong antiviral activity against SARS-CoV-2. The antiviral results performed with the phage phi 6 are shown in Figure 3.

**Figure 3.**
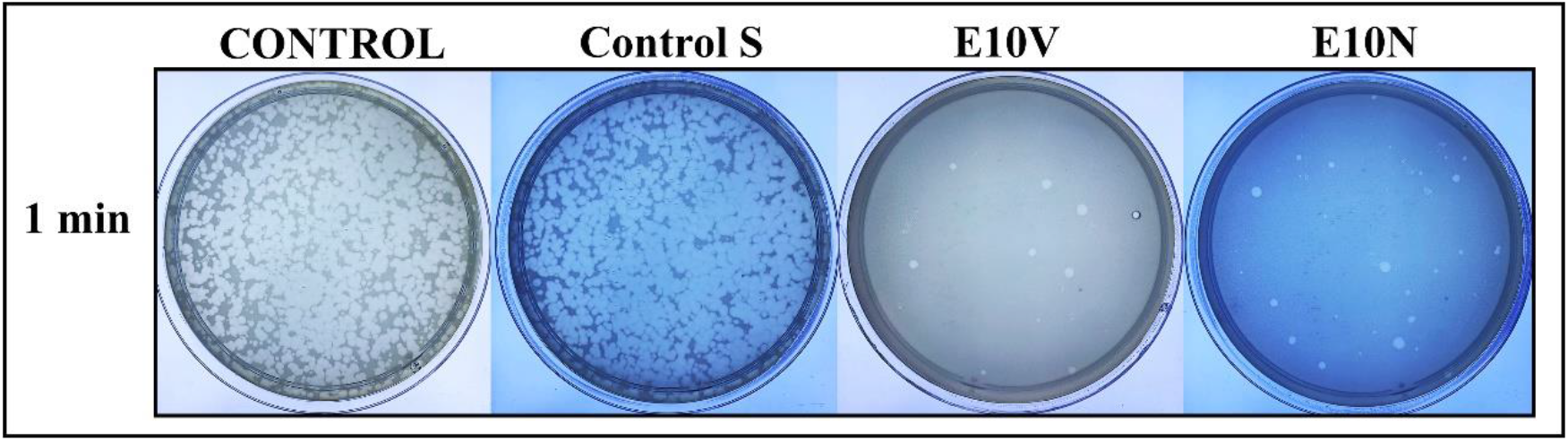
Phage phi 6 viability determined by the double-layer method. Titration images of undiluted samples for control, untreated non-woven fabric (Control S), and cranberry extract samples with biofunctional coating of cranberry extract of VITAFAIR (E10V) or NUTRIBIOLITE (E10N) at 1 minute of viral contact.

Thus, the phage phi 6 losses infectivity after being in contact with both types of smart fabrics for 1 minute and thus very few plaques are observed on the bacterial lawns. However, a similar amount of plaques is observed in the control and untreated fabric on the bacterial lawns after the same time of contact (see Figure 3). The phage titers in PFU/ml of each type of sample were calculated and compared with the control in log reductions (see Figure 4).

**Figure 4.**
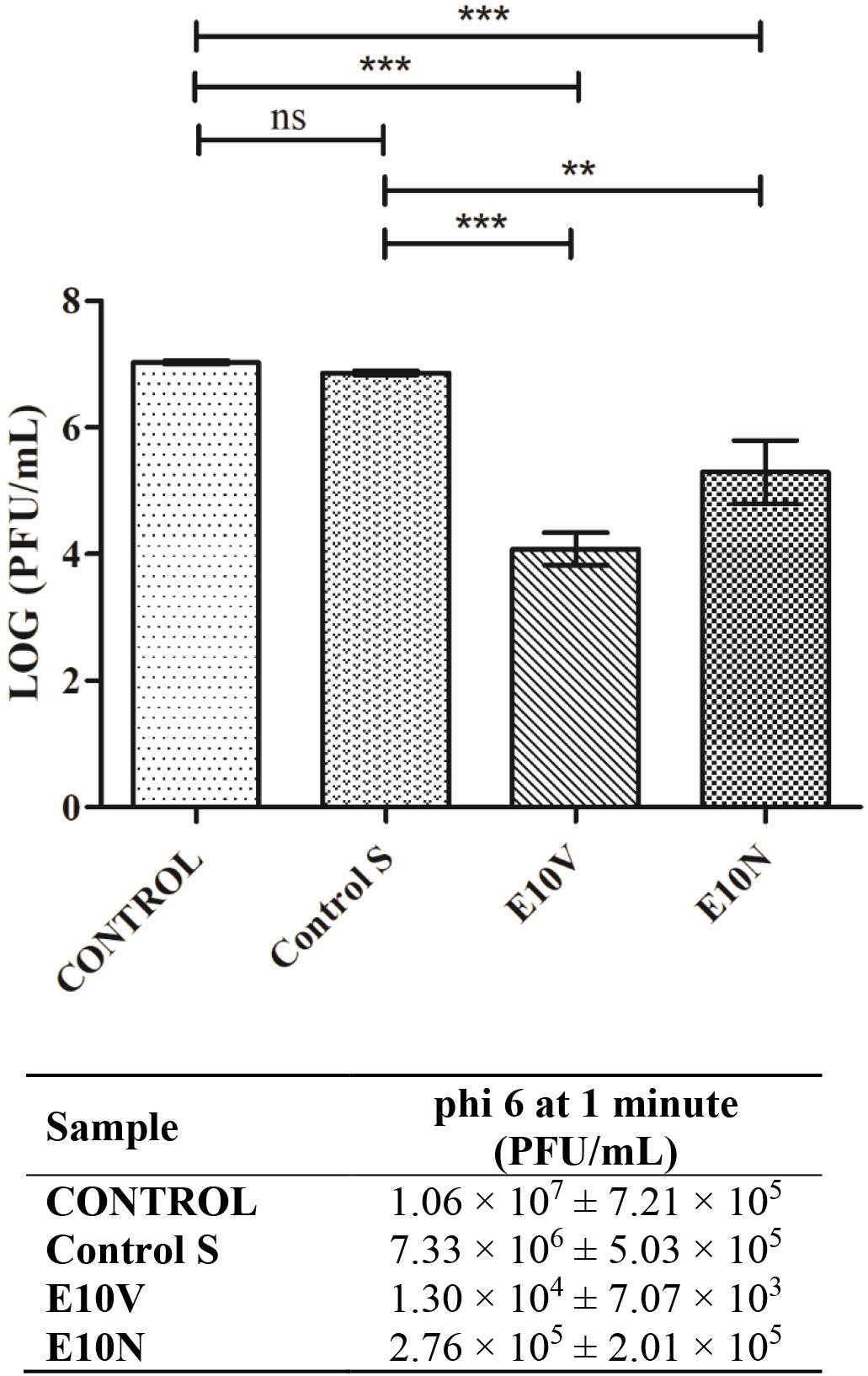
Reduction of infection titers of the phage phi 6 in logarithm of plaque-forming units per mL (log(PFU/mL)) and in PFU/mL measured by the double-layer method. Untreated non-woven fabric (Control S), and non-woven fabric with biofunctional coating of cranberry extract of VITAFAIR (E10V) or NUTRIBIOLITE (E10N) at 1 minute of viral contact. ^***^ *p* > 0.001; ^**^ *p* > 0.01; ns, not significant.

Figure 4 shows that the titers obtained by contacting the phages with the Control S for 1 minute are similar to the CONTROL. However, the phages in contact with the fabrics containing cranberry extract (E10V and E10N) for 1 minute displayed a statistically significant phage infectivity reduction.

Therefore, these results of SARS-CoV-2 and the phage phi 6 clearly show the potent antiviral activity of the cranberry extracts independently of the commercial brand confirming our first hypothesis of antiviral activity.

### 3.2. Antibacterial results

The antibacterial results obtained by the disc diffusion test are shown in Figure 5.

**Figure 5.**
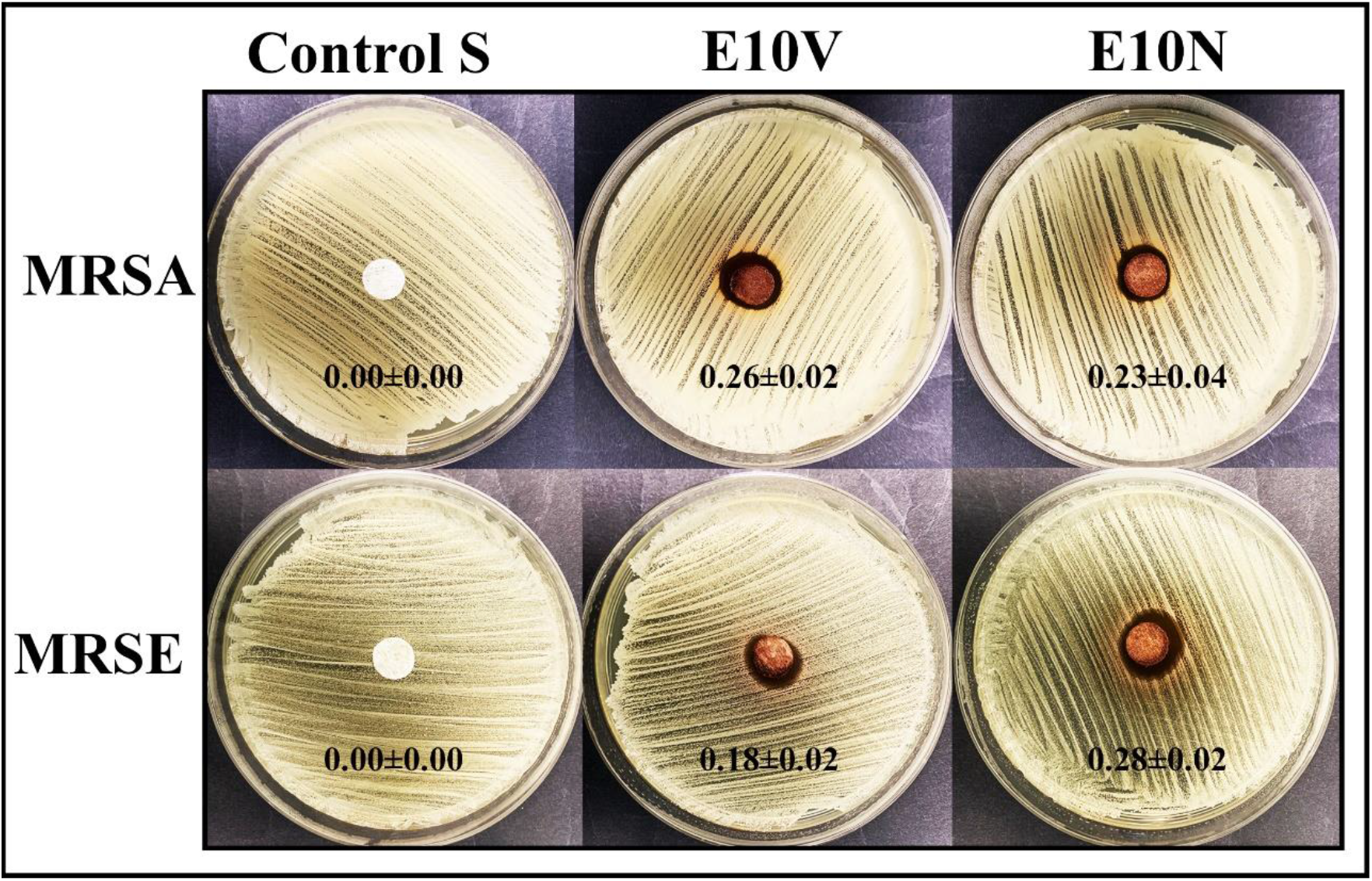
Agar disc diffusion tests. Control sample (Control S), and the fabric with the two cranberry extracts: VITAFAIR (E10V) and NUTRIBIOLITE (E10N). Aerobic incubation for 24 hours at 37 °C. The normalized widths of the antibacterial halos (*nw*_*halo*_) calculated with equation (1) are shown as mean±standard on each image.

Therefore, both non-woven fabrics with biofunctional coatings of cranberry extract showed potent antibacterial properties against methicillin-resistant bacteria *Staphylococcus aureus* and *Staphylococcus epidermidis* confirming our second hypothesis of antibacterial activity.

### 3.3. *In vivo* toxicity

*In vivo* toxicity was studied with the *C. elegans* model, which is a nematode that can be handle at low cost using standard *in vitro* techniques ^42^. Furthermore, it presents fewer ethical problems and shares many genes and signaling pathways with humans. Unlike cytotoxicity assays, *C. elegans* toxicity tests provide data from a whole animal with intact and metabolically active digestive, reproductive, endocrine, sensory and neuromuscular systems^43^. Thus, survival rate, growth and reproduction after an exposure of 24 hours was analyzed with the extracts of the non-woven fabrics with coating of cranberry extracts (see Figure 6).

**Figure 6.**
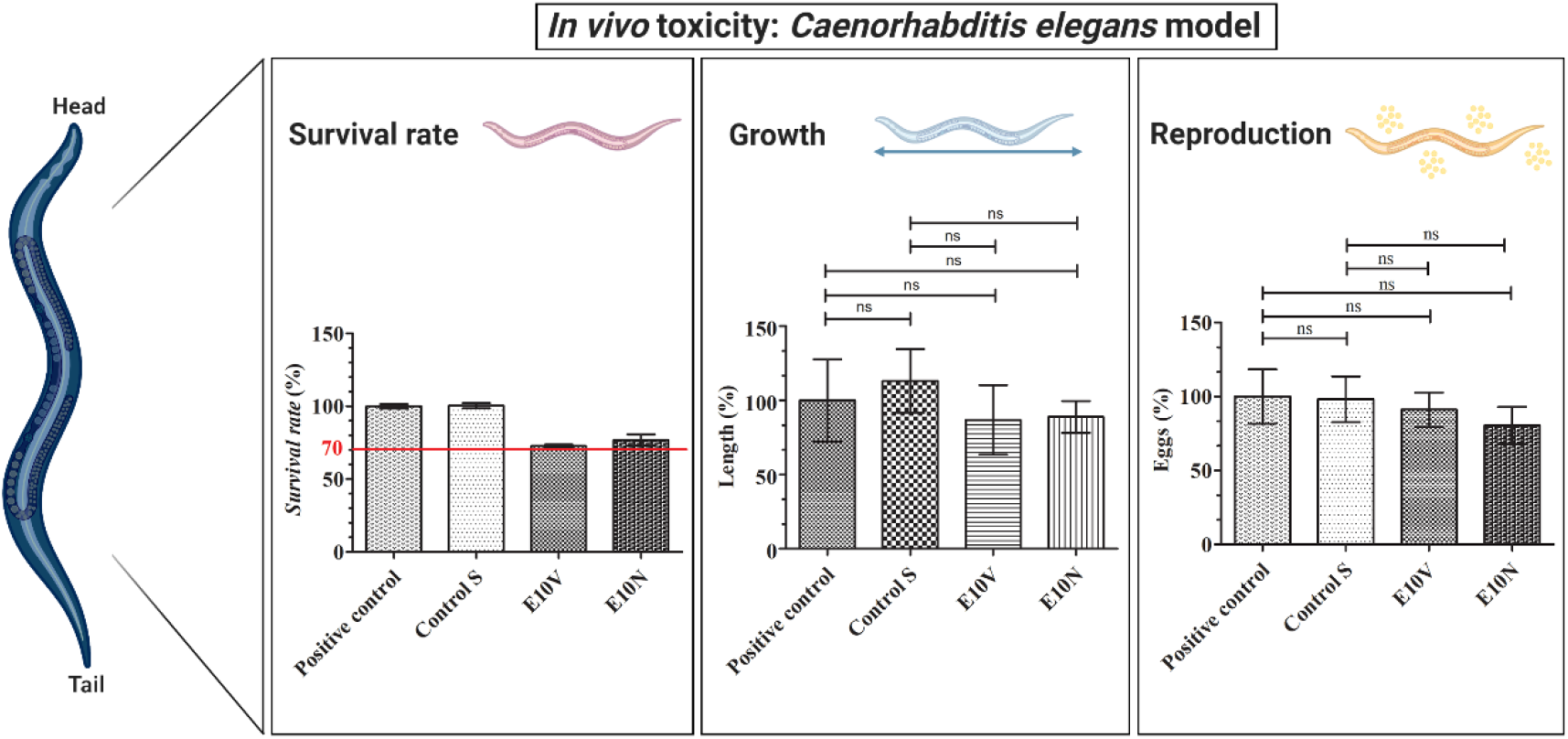
*In vivo* toxicity in *Caenorhabditis elegans* model: survival rate, growth and reproduction after exposure with the extracts of the untreated non-woven fabric (Control S), the extracts of the non-woven fabric treated with VITAFAIR (E10V) and the extracts of the non-woven fabric treated with NUTRIBIOLITE (E10N) with respect to the positive control (100%). Results are shown as mean±standard.; ns, not significant

Both smart fabrics (E10V and E10N) showed a survival reduction lower than 30% in comparison with *C*.*elegans* in the optimal growth conditions (positive control). Furthermore, the extracts of the E10V and E10V non-woven fabrics did not show any effect on growth or reproduction showing the nematodes similar length and number of eggs after 24 hours of exposure. Therefore, these composite materials can be considered non-toxic, especially for infection prevention clothing applications where the materials are used outside the body. It is of note that the antimicrobial non-woven fabrics developed in this study have been produced with cranberries, which are natural and biodegradable products that can be easily grown and thus provide great promise in the fight against SARS-CoV-2. Thus, the fabrication procedure presented here can be used to produce many types of face masks and other next-generation infection prevention clothing such as caps, scrubs, shirts, trousers, disposable gowns, overalls, hoods, aprons and shoe covers to inactivate enveloped viruses such as SARS-CoV-2 (see Figure 7).

**Figure 7.**
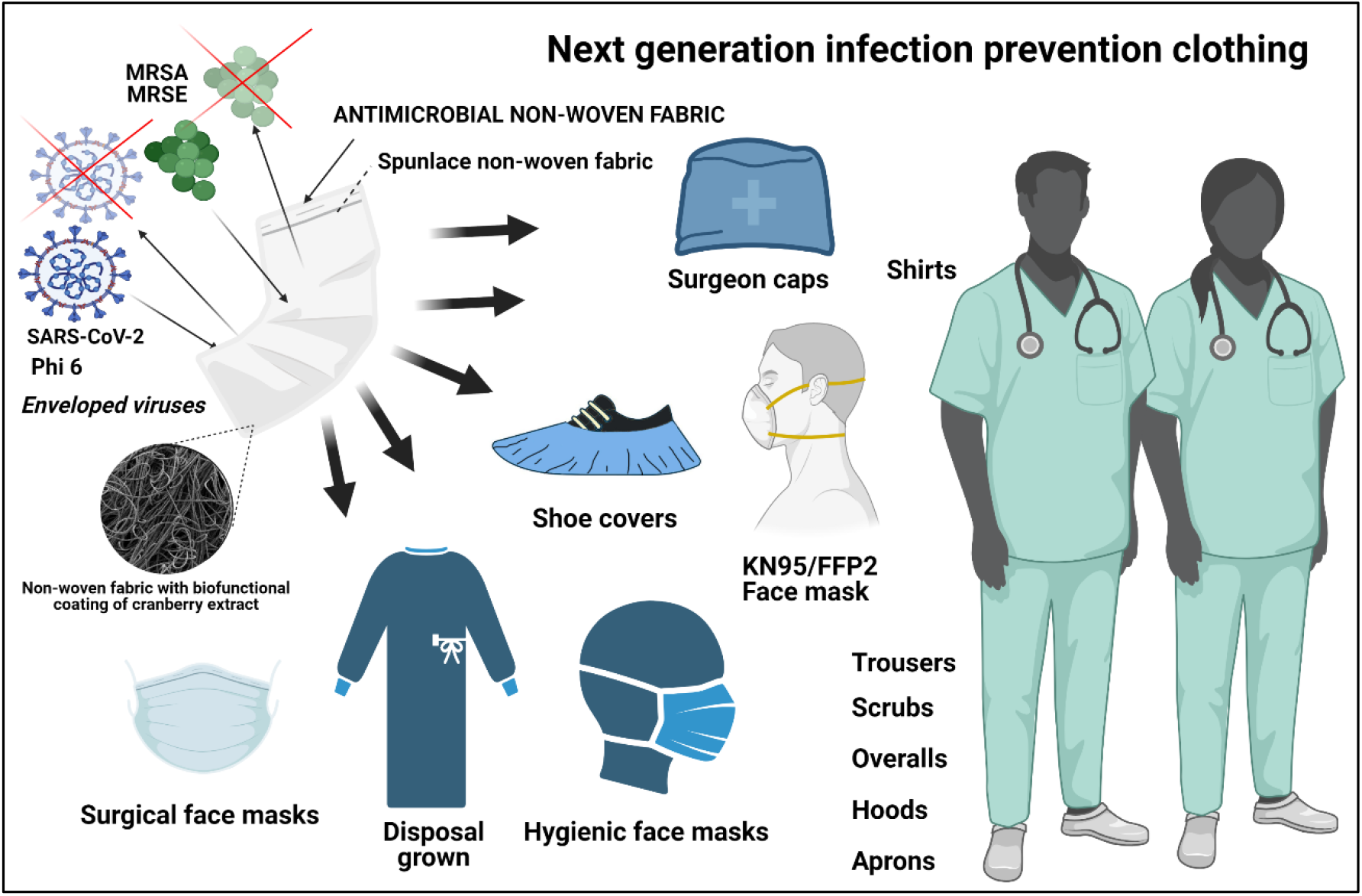
Antiviral infection prevention clothing produced with the developed antiviral low-cost technology capable of inactivating enveloped viruses such as SARS-CoV-2 and phi 6, and the methicillin-resistant bacteria *Staphylococcus aureus* (MRSA) and methicillin-resistant *Staphylococcus epidermidis* (MRSE) bacteria.

## 4. Conclusion

Two different non-woven fabrics have been developed with two types of commercial cranberry extracts by dip coating. Both composite fabrics showed no toxicity in a *Caenorhabditis elegans in vivo* model and high antiviral activity against both enveloped viruses SARS-CoV-2 and phage phi 6 after just one minute of contact. Since cranberry extracts have also exhibit antiviral activity against other enveloped viruses such as HSV-1, HSV-2, and IFV, the idea that PACs produce strong alterations of their envelope glycoproteins achieving their inactivation is becoming more and more consistent. Furthermore, the non-woven antiviral fabrics showed potent antimicrobial activity against the Gram-positive methicillin-resistant bacteria *Staphylococcus aureus* and *Staphylococcus epidermidis*. Therefore, further research could elucidate the natural and biodegradable broad-spectrum treatments for the next generation of infection prevention clothing in the current COVID-19 pandemic.

## Acknowledgments

The authors would like to express their gratitude to the Fundación Universidad Católica de Valencia San Vicente Mártir for their financial support through the Grant 2020-231-006UCV and to the Ministerio de Ciencia e Innovación (PID2020-119333RB-I00 / AEI / 10.13039/501100011033) (awarded to Á.S-A). This research was also supported by grants from the Japan Agency for Medical Research and Development (AMED) (20fk0108270h0001, 20fk0108263s0201). This work was supported by Joint Usage/Research Center program of Institute for Frontier Life and Medical Sciences Kyoto University. We would like to thank Dr. Yoshio Koyanagi and Dr. Kazuya Shimura (Kyoto University) for setup and operation of the BSL-3 laboratory.

## Author Contributions

Conceptualization, methodology, validation and formal analysis: M.M., K.T. and Á.S.-A.; software: K.T. and Á.S-A.; investigation: A.T-M., A.C-V., M.M., Y.M., T.N., K.T. and Á.S-A.; resources: M.M., K.T. and Á.S-A.; data curation: A.T-M., A.C-V., K.T. and Á.S-A.; visualization: K.T. and Á.S-A.; writing—original draft preparation: Á.S-A.; writing—review and editing: A.T-M., A.C-V., M.M., K.T. and Á.S-A.; supervision: M.M., K.T. and Á.S-A.; project administration: K.T. and Á.S-A.; funding acquisition: K.T. and Á.S-A. All authors have read and agreed to the published version of the manuscript.

